# Widefield light sheet microscopy using an Airy beam combined with deep-learning super-resolution

**DOI:** 10.1101/2020.02.27.967547

**Authors:** Stella Corsetti, Philip Wijesinghe, Persephone B. Poulton, Shuzo Sakata, Khushi Vyas, C. Simon Herrington, Jonathan Nylk, Federico Gasparoli, Kishan Dholakia

**Affiliations:** SUPA, School of Physics and Astronomy, University of St Andrews, North Haugh, St Andrews, Fife, KY16 9SS, UK; Strathclyde Institute of Pharmacy and Biomedical Sciences, University of Strathclyde, Glasgow, G4 0RE, UK; Hamlyn Centre for Robotic Surgery, Imperial College London, South Kensington Campus, London, SW7 2AZ, UK; Nicola Murray Centre for Ovarian Cancer Research and Edinburgh Pathology, Cancer Research UK Edinburgh Centre, MRC Institute of Genetics and Molecular Medicine, University of Edinburgh, EH4 2XR, UK; Department of Physics, College of Science, Yonsei University, Seoul 03722, South Korea

## Abstract

Imaging across length scales and in depth has been an important pursuit of widefield optical imaging. This promises to reveal fine cellular detail within a widefield snapshot of a tissue sample. Current advances often sacrifice resolution through selective sub-sampling to provide a wide field of view in a reasonable time scale. We demonstrate a new avenue for recovering high-resolution images from sub-sampled data in light-sheet microscopy using deep-learning super-resolution. We combine this with the use of a widefield Airy beam to achieve high-resolution imaging over extended fields of view and depths. We characterise our method on fluorescent beads as test targets. We then demonstrate improvements in imaging amyloid plaques in a cleared brain from a mouse model of Alzheimer’s disease, and in excised healthy and cancerous colon and breast tissues. This development can be widely applied in all forms of light sheet microscopy to provide a two-fold increase in the dynamic range of the imaged length scale. It has the potential to provide further insight into neuroscience, developmental biology and histopathology.

## 1 Introduction

Widefield optical imaging at depth has opened up new vistas for neuroscience and developmental biology [1, 2, 3], and is expanding its remit into histopathology [4]. There, the capacity to image across length scales is an important pursuit, seeking to maximise the field of view, whilst capturing the fine details needed to discern cellular-scale morphology in three dimensions [5]. Several non-destructive depth-sectioning fluorescence microscopy techniques, such as confocal microscopy [6] and non-linear microscopy [7] have been pioneering in these areas. Recent advances in high-resolution imaging with structured illumination microscopy (SIM) [8,9] or at ultraviolet excitation wavelengths (MUSE) [10] have revealed novel information on the sub-cellular scale; however, they have often been at the sacrifice of the size of the area probed, expense or speed.

An emerging candidate in this area is light sheet microscopy (LSM) [5]. In particular, the geometry of LSM, with orthogonal illumination and detection arms, has shown a significant reduction in photodamage accompanied with rapid widefield volumetric image acquisition. This has been married with new photonics innovations, such as the use of propagation-invariant light fields, and multi-objective schemes, offering a multiview sample perspective and a high isotropic resolution [11, 12]. Further advances have also taken place in sample administration and orientation [13, 4, 14]. To address the notion of gaining information at differing scales simultaneously, super-resolution has been coupled with the geometry of the LSM through the approach of using structured illumination [15] or localisation [16, 17].

A more facile approach for imaging across scales would be to employ recent innovations in deep learning [18] to retain the wide field of view afforded by light sheet microscopy and to computationally recover a super-resolved image. Whilst a high optical resolution can be accomplished with the use of structured light fields that can carry extended spatial frequency information, many widefield systems suffer from the need to Nyquist sample the field of view, which fundamentally limits the dynamic range of the spatial length scale. In this paper, we breach this limit to resolution with the use of deep learning to recover high-resolution images from sub-sampled information. We achieve this by using a generative adversarial network (GAN) model pre-trained on widefield fluorescence images [18]. We couple this with the particular use of a propagation-invariant Airy light field for illumination, which allows for a wider field of view captured on the camera than when using a Gaussian beam; furthermore, it has inherent advantages in achieving increased depth penetration through aberrating tissues [19]. In this way, we realise imaging across length scales with LSM over extended fields of view and depth ranges.

Our embodiment is an open-top geometry that is inspired by and builds upon the LSM system developed by Glaser et al. [4]. In particular, we also make use of a solid immersion lens (SIL) placed underneath the sample plane. A SIL is a high refractive index glass hemisphere that increases both the illumination and detection numerical aperture by its refractive index. Compared to the use of a high-magnification objective, it has the advantage of providing a high resolution while retaining a wide lateral field of view (along the sample plane). The orthogonal illumination and detection arms of the LSM are aligned through the same SIL, tilted at 45° the sample plane. In this geometry, the field of view in depth is effectively set by the depth of focus of the light sheet. Consequently, using a conventional Gaussian light sheet inherently limits the depth range. We overcome this limit by using a propagation-invariant Airy light sheet to preserve a wide field of view over the entire imaging volume. Our use of both the Airy and the Gaussian light sheets with the SIL enables us to capture 3-mm lateral fields of view. However, the use of the Airy light sheet enables us to capture a 1.6-mm depth range in one shot, compared to a 60 *μm* depth range when using a Gaussian light sheet with the same numerical aperture. Furthermore, the combined use of deep learning with the Airy light sheet leads to a two-fold improvement in the spatial resolution and an improvement to image contrast. We characterise these improvements on test phantoms with fluorescent beads. Further, we demonstrate the performance of our system on excised brain tissue from a mouse model of Alzheimer’s disease, revealing regions of plaque, as well as on unfixed excised human colorectal and breast tumour samples. We demonstrate the capability of our setup to image turbid tissues in depth maintaining a good resolution over a wide field of view. This framework may be broadly applied to other LSM geometries without the need for training a bespoke neural network.

## 2 Materials and Methods

### 2.1 Optical setup

The LSM setup features an open-top geometry enabled by a solid immersion lens [4]. We introduce several additional key capabilities, namely, a variable Gaussian or Airy beam illumination profile, a wider field of view, a zoom lens in the detection and novel image processing methods based on deep learning. Figure 1 illustrates the sample stage and the image acquisition procedure. The system employs a 488-nm wavelength laser (STRADUS-488-150, Vortran). The illumination power was kept below240 *μ*W at the sample, modulated using a waveplate and a polarisation beam splitter. After expanding the beam, the illumination is split into two parallel optical paths, with one path modulated by a one-dimensional cubic polynomial phase mask (PM) forming an Airy beam, such that the Airy or the Gaussian illumination profile can be selected during imaging by using the shutters. The shape and properties of the Airy beam are dictated by the strength of the cubic phase modulation. The phase modulation profile of the phase mask is described by:

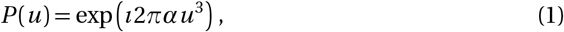

where *u* is the normalised pupil coordinate normal to the imaging plane and *α* is a dimensionless parameter that sets the propagation invariance of the Airy beam [11]. In particular, the propagation-invariant length increases with *α*, with *α* = 0 being equivalent to a Gaussian beam. The value of *α* used was estimated to be 5 from the beam profile. For the equivalent beam waist, the theoretical depth of focus is 60 *μ*m for the Gaussian beam and 1.6 mm for the Airy beam. Due to the tilted geometry (Fig 1), the depth and the light sheet propagation are not coaxial. For consistency, we term the lateral and depth imaging ranges as the field of view.

**Figure 1:**
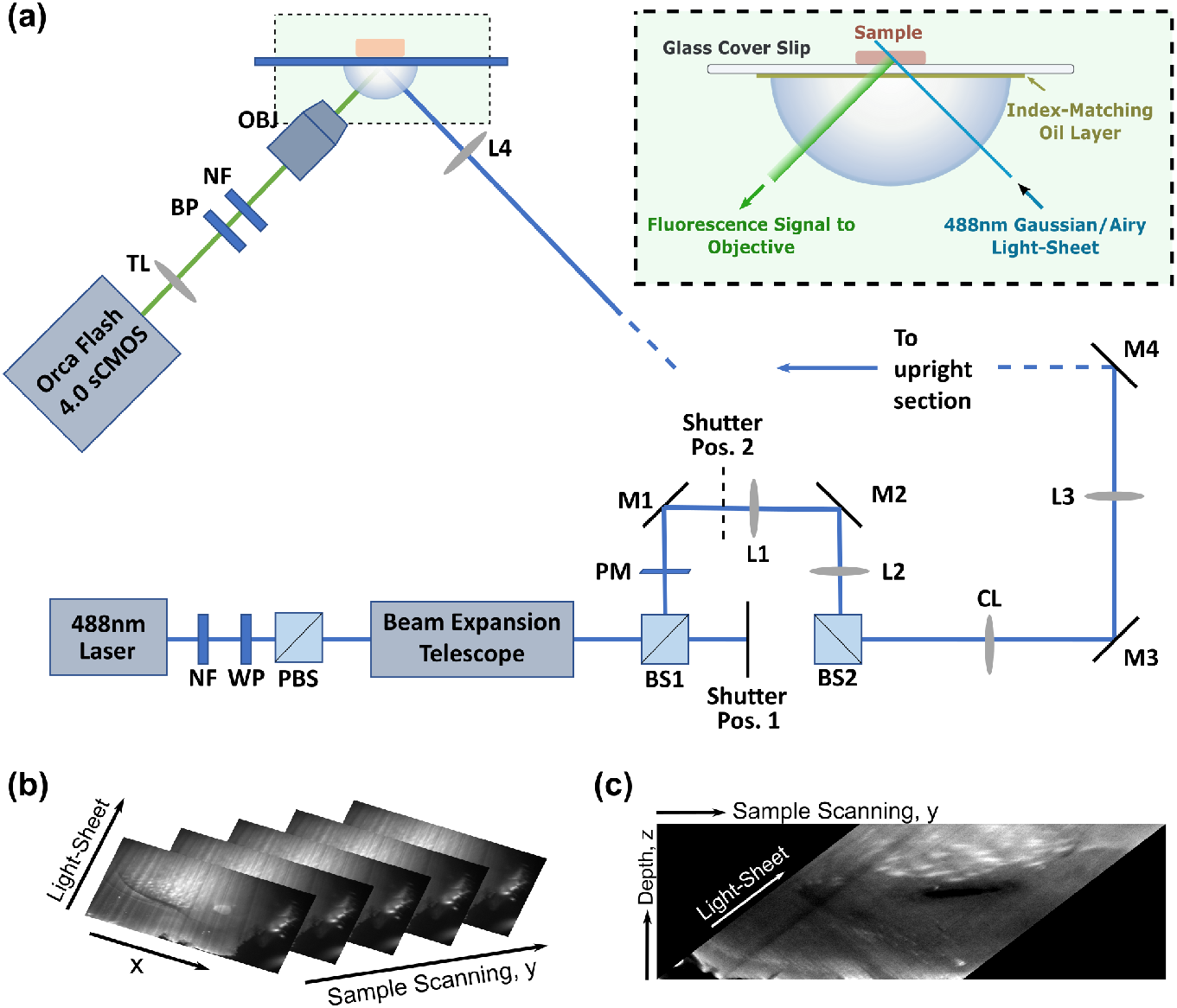
Setup and acquisition process. (a) Schematic of the setup. NF: notch filter; WP: waveplate; PBS: polarising beam splitter; BS1, BS2: 50:50 beam splitters; PM: Airy phase mask; M1-4: mirrors; L1-4: lenses; CL: cylindrical lens; OBJ: objective; BP: bandpass filter; TL: tube lens. In the inset: A solid immersion lens (SIL) and an index-matching oil layer are used for imaging a sample on top of a glass coverslip. The sample is translated with the coverslip and (b) a series of images are recorded. (c) The image sequence is interpolated into the physical coordinate space.

The Gaussian and Airy paths are recombined and reshaped by a cylindrical lens (CL) into a light sheet and focused into the sample, one modality at a time, through a 15-mm diameter fused-silica solid immersion lens (SIL, BMV optical, Ottawa, ON Canada), a thin layer of immersion oil and a 1-mm thick glass coverslip to reach the sample plane (Fig. 1(a)). The SIL provides a magnification factor that is equivalent to its refractive index (n = 1.46) giving an illumination numerical aperture (NA) of 0.1. The SIL reduces aberrations from refractive index mismatch resulting from the illumination passing off-axis to the cover slide [4]. This enables the sample to be placed and imaged on top of the cover slip without additional mounting or embedding.

The fluorescence from the sample is collected by a detection objective (OBJ: MRD00045FL50 mm, 4XNA0.2, CFI Plan Apochromat Lambda, Nikon), then is passed through an achromatic tube lens (TL: AC254-150-A-ML FL 150 mm, Thorlabs) and focused onto the sCMOS camera (ORCA-Flash4.0, Hamamatsu) after passing through a 488-nm notch filter (NF) and a 532-nm band pass filter (BP: 50-nm bandwidth). The total magnification of the collection arm, including the SIL, is 4.3, the NA is 0.29, and the camera field of view is 3 mm. Additionally, we introduce a zoom lens (MVL6X12Z, Thorlabs) after the tube lens to provide an adaptable field of view and magnification. The zoom lens allows higher sampling resolution by reducing the field of view down to a minimum of 675 *μ*m.

A custom-made sample stage is attached to a motorised translation stage (M-230.10 with a C-863.11 DC motor controller, Physik Instrumente). The stage enables sample scanning at 45 degrees to the detection axis (Fig. 1(b)), such that a 3D image stack can be acquired. The whole setup fits upon a small breadboard (dimensions 35×50×60 cm).

### 2.2 Image acquisition and processing

The imaging plane was oriented at 45 degrees to the sample translation. During imaging, the sample was translated and a sequence of images was recorded (Fig. 1(b)). Custom Matlab software was developed to interpolate the image sequence into the physical coordinate space (Fig. 1(c)). For an image stack represented by a matrix M with pixel coordinates (*n, m, l*), a translation distance *δy* and a camera pixel size of *δx*, the transformation to the physical space becomes: 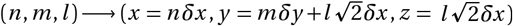, where *x* is the transverse light sheet direction, *y* is the sample scanning vector and *z* is the depth from the cover slip, with (*x, y, z*) being orthogonal coordinates.

Images taken with the Airy illumination are further deconvolved using a twodimensional point-spread function (PSF) calculated from theory [11]. Briefly, the PSF is generated from a Gaussian intensity and an Airy phase modulation function given in Eq. (1). This is done efficiently by taking the partial spectrum components in the spatial domain using the chirp z-transform combined with the defocus function set by the objective NA, the entrance pupil and the mean refractive index of the sample. The transform is sampled such that it has the same spatial pixel size as the recorded LSM data. The Airy illumination profile is further apodised by the Gaussian detection, reconstructing the final theoretical two-dimensional PSF.

Deconvolution is performed in the *zy* plane (depth, scanning plane) using the Richardson-Lucy algorithm [20] and repeated for all positions in *x*. In an ideal system, the detected image, *D*, is given by the convolution of the sample intensity, *I*, and the PSF with some added noise 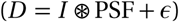. The deconvolution algorithm recovers an estimate, 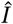, by iterating:

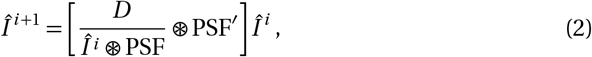

where PSF’ is an inverted PSF, *i.e.,* PSF’(**r**) = PSF(-**r**). The estimate ofthe algorithm is the maximum-likelihood solution based on Bayes’ theorem in the presence of noise [20].

Following deconvolution, deep-learning super-resolution was performed based on the GAN model architecture originally presented in Wang et al. [18]. In that work, they demonstrated a widely applicable improvement in resolution across various imaging modalities, including epifluorescence, TIRF and STED microscopies. The model in this paper is based on the pre-trained set that maps low resolution (10x, 0.4 NA) images to higher-resolution co-registered (20x, 0.75NA) images in widefield epifluorescence microscopy. We found this approach to be well-suited for the LSM system, for both the Airy and Gaussian illuminations, as we detail in the Results and Discussion sections. It breaches a major hurdle to the broad adoption of the technique, which is the hardware-limited and computationally demanding need to train a network for each individual imaging system. This algorithm employs a GAN architecture to achieve super-resolution from a single-image input [21]. The architecture and validation are described in detail in [18], and the pre-trained model is openly available. Briefly, training was done by pixel-wise mapping of a set of high and low resolution images with sub-pixel-accurate co-registration. The algorithm then creates an estimate of the high resolution image through a generative network, whilst simultaneously a discriminative network is trained to blindly discriminate between the estimates and the equivalent co-located high-resolution images. The process is back-propagated for the high-resolution image mapping to the low-resolution domain. An adversarial loss regularised by a mean-squared error and structured-similarity estimators [21, 18] are used to converge iterations of this process, such that the high-resolution estimates from the low-resolution images are indistinguishable from the high-resolution ground truths. This metric of effectiveness is evaluated on a subset of test images that are not used in the training of the network.

Custom-built Matlab software is used to interface with the deep-learning module developed in [18]. The LSM images are decomposed into a sequence of 1024×1024-pixel frames, splitting individual cross-sections into a series of mosaics. The frames are sequentially run through the network and are re-stitched back into the original orientation. An overlapping margin exceeding the maximum network convolution size is used to prevent stitching artefacts along image boundaries. This process takes several seconds per *xy* -frame when run on the CPU, with a substantial speed-up expected with a GPU implementation.

### 2.3 Sample preparation and staining

#### 2.3.1 Characterisation of phantom

200 nm diameter green fluorescent microspheres were mixed with 1.5% agarose and used for characterization of the system’s PSF. 4.8 *μ*m diameter green fluorescent microsphere were mixed with 1.5% agarose and used for the calibrating the setup with objects of known shape and size. The samples were pipetted on a glass coverslip and then placed on top of the setup.

#### 2.3.2 Cleared mouse brain

All experimental procedures were performed in accordance with the United Kingdom Animals (Scientific Procedures) Act of 1986 Home Office regulations and approved by the Home Office (PPL 70/8883). A 5xFAD::ChAT-IRES-Cre mouse (male, 3 months old) (5xFAD, JAX006554) (ChAT-IRES-Cre, JAX006410) was deeply anesthetized with a mixture of pentobarbital and lidocaine and perfused transcardinally with 20 mL saline followed by 20 mL4 % paraformaldehyde/0.1 M phosphate buffer, pH 7.4. The brain was removed and immersed in the same fixative solution overnight and kept in phosphate buffered saline (PBS) with 0.02 % sodium azide at 4^°^C.

The brain tissue was trimmed into half hemispheres and processed with thioflavinS (0.05%). The tissue was pre-treated, stained and cleared following the method described in Renier et al. [22,23]. First, the tissue was dehydrated with a gradual addition of methanol in distilled water (20%, 40%, 60%, 80%, 100%, each for 1 hour) at room temperature. Then the tissue was bleached with 5% hydrogen peroxide in ice-cold methanol at 4^°^C overnight. After washing with 100% methanol at room temperature (twice, each for 1 hour), the tissue was rehydrated by gradual removal of methanol in distilled water (80%, 60%, 40%, 20%, each for 1 hour), followed by immersion in PBS for 1 hour. After washing the tissue with 0.2% Triton X-100 in PBS (twice, each for 1 hour) at room temperature, the tissue was incubated in a permeabilization solution (0.2% Triton X-100, 0.3 M glycine and 20% DMSO in PBS) at 37°C for two days. Then the tissue was incubated in a staining solution (0.05% thioflavin-S, 0.2% Triton X-100, and 10% DMSO in PBS) at 37^°^C for three days.

After washing with 0.2% TritonX in PBS (four times, each for 1 hour) at room temperature, tissue was left in the same solution overnight in the dark. The tissue was then gradually dehydrated again (20-100% methanol, each for 1 hour) at room temperature, followed by clearing the tissue with 66% dichloromethane (DCM) in methanol at room temperature overnight. After washing with 100% DCM for 15 min twice, tissue was incubated in 100% dibenzyl ether overnight. Tissue was kept in the same solution at 4°C until imaged.

#### 2.3.3 Colorectal tissues

Matched colorectal adenocarcinoma and normal tissue samples were provided frozen by the Tayside Tissue Bank, Ninewells Hospital and Medical School, Dundee (Tissue request and approval no. TR000289). Each sample of approximately 1×0.5×0.7 cm in size was thawed in phosphate-buffered saline (PBS) at ambient temperature for 5 minutes. The defrosted tissues were then stained for 40 seconds with 1 mM of acridine orange and then washed for 10 seconds in PBS. They were then blotted dry and immediately imaged with the system.

#### 2.3.4 Breast tissues

Breast tissue specimens were obtained from consented patients undergoing breast surgery at Charing Cross Hospital in London using the Imperial tissue bank license (Project R12047). Small cut-outs measuring approximately 5×5 mm across were obtained each from the tumor and fibrofatty tissues distant from the tumor site. The samples were snap frozen at −80°C immediately following surgery. The samples were later thawed at an ambient temperature for 5 minutes, stained with 0.02 % acriflavine hydrochloride solution for 1 minute, rinsed with PBS for 30 seconds and imaged immediately.

## 3 Results

### 3.1 Deeper imaging with an Airy light field

We experimentally compare the effective field of view at depth of the Gaussian and Airy light sheets by using a test phantom comprising sub-resolution 200-nm isolated green fluorescent microspheres embedded in 1.5% agarose (sample refractive index ~ 1.33). Figures 2(a) and (b) show the maximum intensity projection in the *xy* plane of the data recorded with the Gaussian beam and the Airy beam, respectively. As expected, the Airy beam demonstrates a longer depth of focus at the expense of an axially elongated and asymmetric PSF (Fig. 2(d)). The Airy PSF, however, retains its spatial frequencies in the optical transfer function and, thus, the spatial information about the sample. We recover this information using a two-dimensional Richardson-Lucy deconvolution, as described in the Methods section. Figure 2(c) shows the results of the deconvolution.

**Figure 2:**
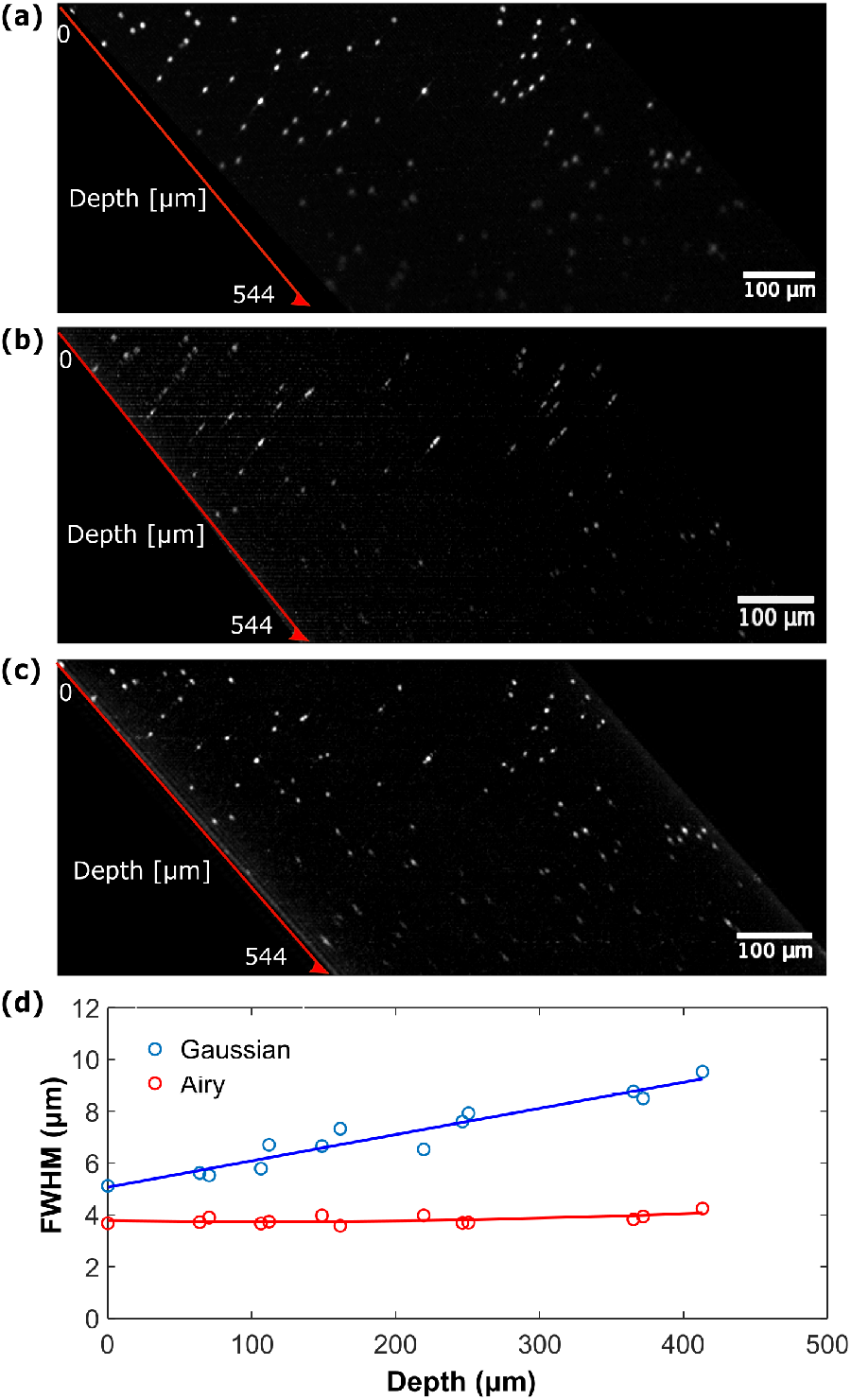
Maximum intensity projections in the *yz* plane of 200-nm blue fluorescent microspheres embedded in 1.5% agarose recorded with the (a) Gaussian light sheet and (b) Airy light sheet. (c) Deconvolution of the experimental data shown in (b). (d) Experimentally determined full-width at half-maximum (FWHM) as function of depth of the sample.

Comparing Figures 2(a) and (c), the Airy beam yields a slight improvement in the penetration depth compared to the Gaussian beam, given as the intensity drop-off of the image in depth. This is because attenuation is set primary by the scattering properties of the sample. However, it is evident that the beads imaged using Gaussian illumination have substantially broader PSFs in depth, normal to the light sheet plane.

We compare this resolving power between the Gaussian and Airy light sheet by the full-width at half-maximum (FWHM) of 13 isolated fluorescent microspheres randomly sampled at different depths in Figs. 2(a) and (c). The FWHM was determined in the axial and lateral directions from line profiles through the beads. The fluorescence detection path is the same for both illumination beams, thus, a similar lateral resolution is expected. The lateral resolution was measured to be 4.7 ± 0.6 *μ*m and 4.2 ± 0.6 *μ*m for the Gaussian and the Airy light sheet, respectively. Figure 2 (d) shows the axial resolution for both Gaussian and Airy light sheet as a function of depth into the sample. It is clear that the PSF of the deconvolved Airy has a substantially finer axial resolution, which is maintained in depth, whilst the Gaussian PSF experiences a substantial defocus. In fact, the flatter curve of the Airy beam demonstrates that it maintains the same focus (axial resolution) over a longer distance when compared to the Gaussian beam. The Gaussian axial resolution was measured to be ~5.3 *μ*m at the surface and then increased with depth to reach the value of 9.5 *μ*m at a distance of400 *μ*m from the surface. The FWHM obtained with the Airy was measured to be ~3.6 *μ*m at the surface and remained quite confined at depth reaching the value of 4.3 *μ*m at 400 *μ*m from the surface. These results agree with what was previously observed by Nylk et al. [19]. The Airy light sheet penetrates deeper into the sample without aberration correction. This may be attributed to a ‘self-healing’ property that allows the wavefront to reconstruct itself after encountering an obstacle [24].

### 3.2 Single-image super-resolution

The minimal lateral resolution of the LSM is set by the diffraction limit and is determined by the numerical aperture of the detection arm. For our system, the theoretical diffraction-limited resolution is 1.1 *μ*m. However, for the maximal camera field of view of 3 mm, the pixel size at the image plane was 1.6 *μ*m. Thus, the Nyquist criterion sets the minimal resolution to 3.2 *μ*m (compared to the experimentally measured 4.2 *μ*m).

To achieve a high spatial resolution over a wide field, we employ a deep-learning algorithm initially demonstrated for single-image super-resolution by Wang et al. [18], and described in the Methods section. To validate this method, we introduce a zoom lens to the detection arm that is capable of providing an on-the-fly adaptable sampling resolution and field of view. First we characterise the resolution of the system by imaging a test phantom comprising sparse 200-nm beads (Figs. 3(a-c)) using Gaussian illumination. Low and high resolution images were acquired with a pixel size of 2 *μ*m and lateral field of view of 4 mm (which was cropped to 2 mm by the zoom lens aperture), and a pixel size of 0.5 *μ*m and a low field of view of 1 mm, respectively. The pixel size of 0.5 *μ*m allows the high resolution image to Nyquist-sample the diffraction limit, which is 1.1 *μ*m. For different magnifications, the pixel size was measured by using a graticule. Figure 3(c) shows the network output of the deep-learning algorithm applied to the widefield low-resolution image. Figures 3(g) and (h) show the vertical and horizontal line profiles through a fluorescent bead marked by the cross target in (a-c). It is clear that the resolution is severely degraded by the low sampling of widefield imaging in the low-resolution case. The FWHM of the line profiles in both the horizontal and vertical directions were measured to be 4.3 *μ*m for the low-resolution image, 2.5 *μ*m for the network output and 1.6 *μ*m for the high-resolution image. The deep-learning algorithm recovers a resolution closer to the high-resolution image but over a widefield. The gain in the field of view is indicated by the grayed-out area in Fig. 3(b).

**Figure 3:**
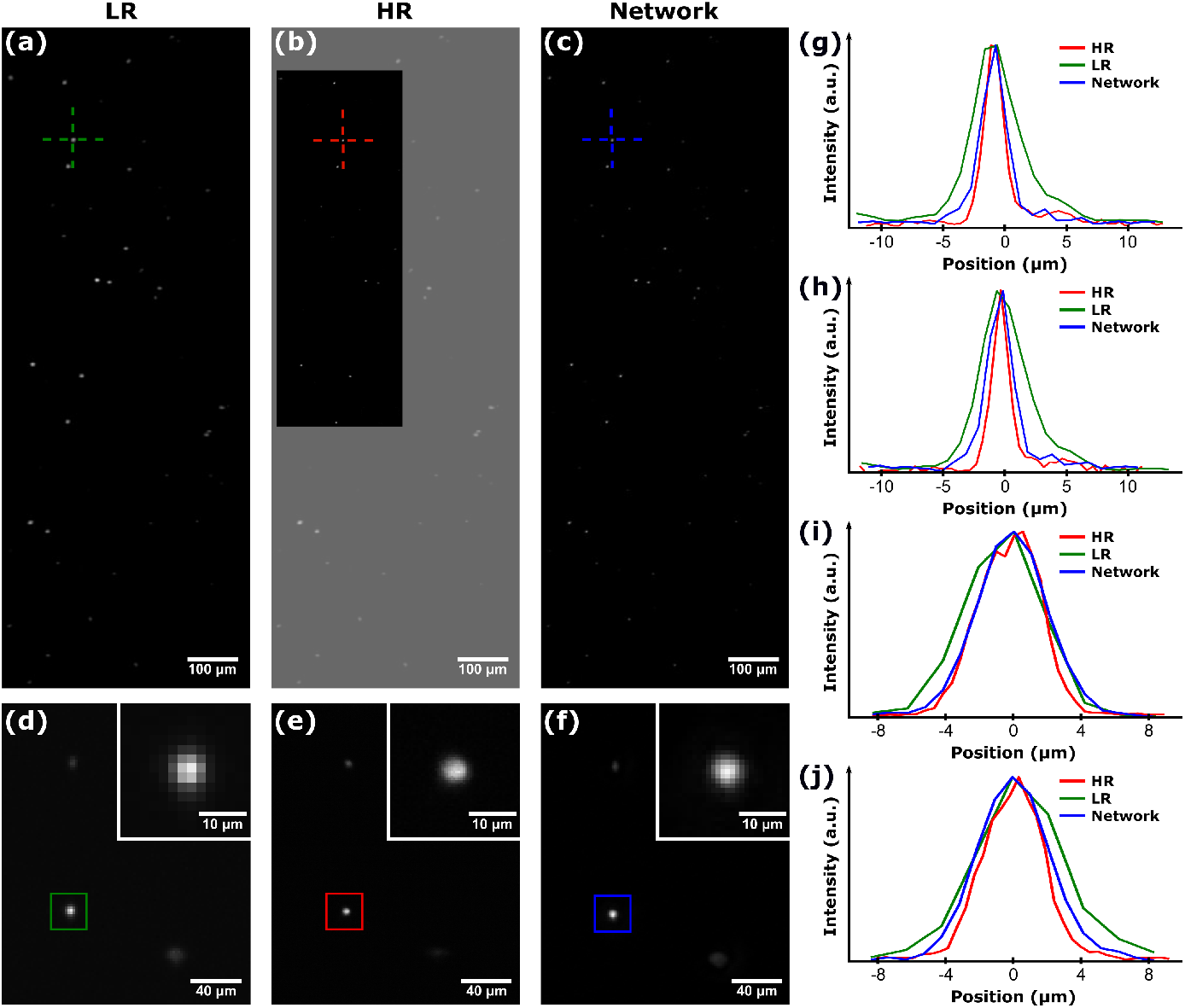
Validation of deep-learning super-resolution on fluorescent beads in agarose. (a-c) 200-nm sub-resolution beads imaged with (a) low resolution (LR), and (b) high resolution (HR) with a use of a zoom lens. In (b), the grayed out area is the LR image, representing the missed field of view. (c) The output of the network applied to the LR image recovers a high resolution over the whole field of view. (g) and (h) shown the vertical and horizontal line profiles across a bead marked by a target. (d-f) 4.8-*μ*m beads imaged with (d) low resolution, (e) high resolution, and (f) output of the network applied to (d). The insets in (d-f) show a magnified view of beads marked by the box. (i) Lateral profile across the bead in (d-f).

We further demonstrate the performance of the deep-learning method on resolvable objects of known size and shape. Figures 3(d-f) show low resolution, high resolution and the network output of 4.8-*μ*m beads imaged with Airy illumination. The vertical and horizontal profiles of the bead marked by the box are presented in Figs. 3(i) and (j), respectively. The FWHM of the vertical line profiles were measured to be 5.4 *μm* for the low-resolution image, 4.6 *μ*m for the high-resolution image and 4.7 *μ*m for the network output. The FWHM of the horizontal line profiles were measured to be 5.8 *μ*m for the low-resolution image, 4.4 *μ*m for the high-resolution image and 4.8 *μ*m for the network output. It is evident that the deep learning method recovers an accurate approximation of the high resolution image, and more closely corresponds to the expected bead size of 4.8 *μ*m.

### 3.3 Cleared mouse brain

To demonstrate the imaging capacity at depth of an Airy light sheet and with deeplearning super-resolution, we imaged a cleared mouse brain from a pre-clinical model of Alzheimer’s disease [25]. The sample comprised a cleared half hemisphere of the brain stained with thioflavin S, as described in the Methods section. The sample was imaged without the zoom lens to achieve the largest 3-mm camera lateral field of view.

Figure 4 shows cross sectional images from the cleared mouse brain at two depths, taken with the Gaussian and Airy light sheets. The Airy data is further deconvolved. The network output of the deep-learning method applied to the Airy images is shown alongside the data. The point-like features that can be observed at a depth of 400 *μ*m, and are highlighted by the yellow squares, are amyloid plaques. The distribution of plaques in the mouse genetic line under study is generally consistent with 5xFAD mice [25]. Amyloid plaques can be confirmed at 3 months old in several brain regions, such as the subiculum, the lateral septum and the neocortex. These plaques are important markers in monitoring the progression of Alzheimer’s disease. In the magnified image of the plaques acquired with the deconvolved Airy light sheet (Fig. 4(e)) compared to the Gaussian light sheet (Fig. 4(d)) the plaques appear sharper. This is to be expected due to the short depth of focus of the Gaussian light sheet compared to the Airy light sheet (see Section 2.1.) After the application of the deep-learning algorithm to the deconvolved Airy image (Fig. 4(c)), the plaques become sharper with a substantial improvement to the image contrast (Fig. 4(f)). The network output provides a further increase in resolution. Since these plaques are substantially larger than the PSF (approximately 10-*μ*m in diameter), the improvement in resolution manifests as an increase in contrast to noise; the resolution and contrast are coupled in an imaging system, and have to be considered in tandem.

**Figure 4:**
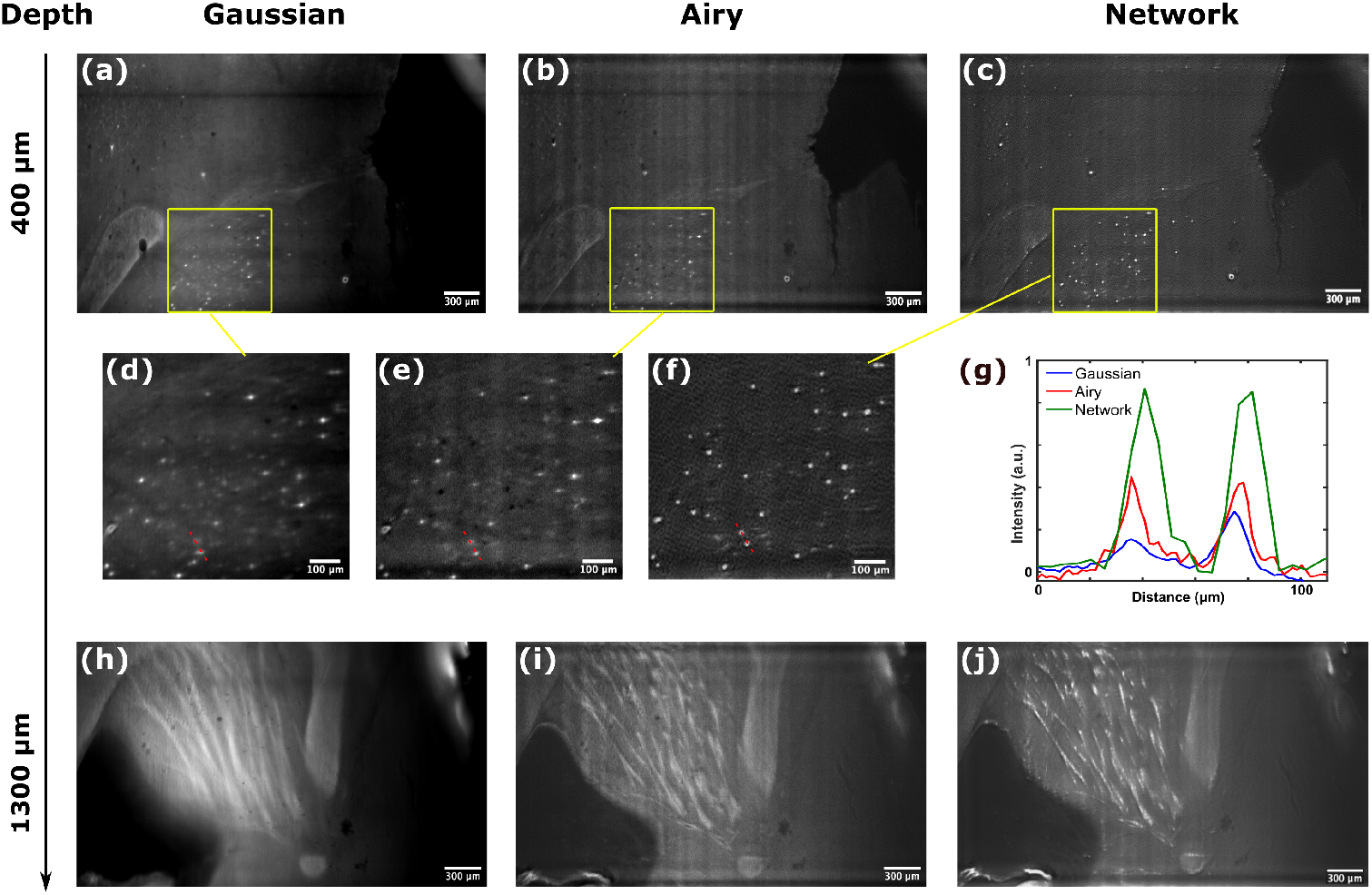
Images of cleared mouse brain tissue stained with thioflavin S. (a-f) Images taken at a depth of 400 *μ*m: (a) Gaussian light sheet, (b) deconvolved Airy light sheet and (c) network outputs of the deconvolved Airy data. (d),(e),(f) are magnified images of the amyloid plaques shown in (a), (b) and (c), respectively. In (d-f) the red dashed lines indicate the (g) line profile through two different plaques. (h-j) Images taken at a depth of 1300 *μ*m: (h) Gaussian light sheet, (i) deconvolved Airy light sheet and (j) network outputs of the deconvolved Airy data.

Figure 4(g) shows a line profile trough two plaques, marked by a red dashed line in Figs. 4(d-f). The line profiles are normalised to the background noise. A better contrast to noise is achieved by the Airy light sheet and improved further by the network. A similar trend is observed from the internal capsule which emitted autofluorescence and was acquired at a depth of 1300 *μ*m (Figs. 4(h-j)). This structure is clearer in the Airy image compared to the Gaussian image, and can be tracked in three dimensions. The capsule is even more evident in the network output of deep learning applied to the deconvolved Airy image (Fig. 4(j)).

### 3.4 Colon tissue

Figure 5 shows images of healthy and cancerous ascending colon tissues taken at various depths with the Gaussian light sheet and with the deconvolved Airy light sheet to which the deep-learning algorithm has been applied. The colon has a smooth surface when viewed macroscopically, but the mucosa, which is imaged here, presents an undulating appearance with occasional infoldings. The surface of the normal colon sample (Fig. 5(a) and (e)) presents numerous colonic crypts. These structures can be followed to a depth of 83 *μ*m in the images taken with the Airy beam, whilst blurring substantially with the Gaussian illumination. Compared to the cleared sample, the depth range in turbid tissue is severely limited by the high scattering. While the advantage of propagation invariance is less evident, the Airy beam has an additional advantage in depth penetration through scattering tissue due to the angular diversity and self healing property of the beam shape [19]. We see this manifest with sharper tissue features and reduced aberrations in depth. The possibility of following structures underneath the surface in three dimensions presents a more facile avenue of assessment to conventional pathology analysis, where tissues are step-sectioned, producing separate 5-*μ*m thick sections that are observed on separate microscope slides.

**Figure 5:**
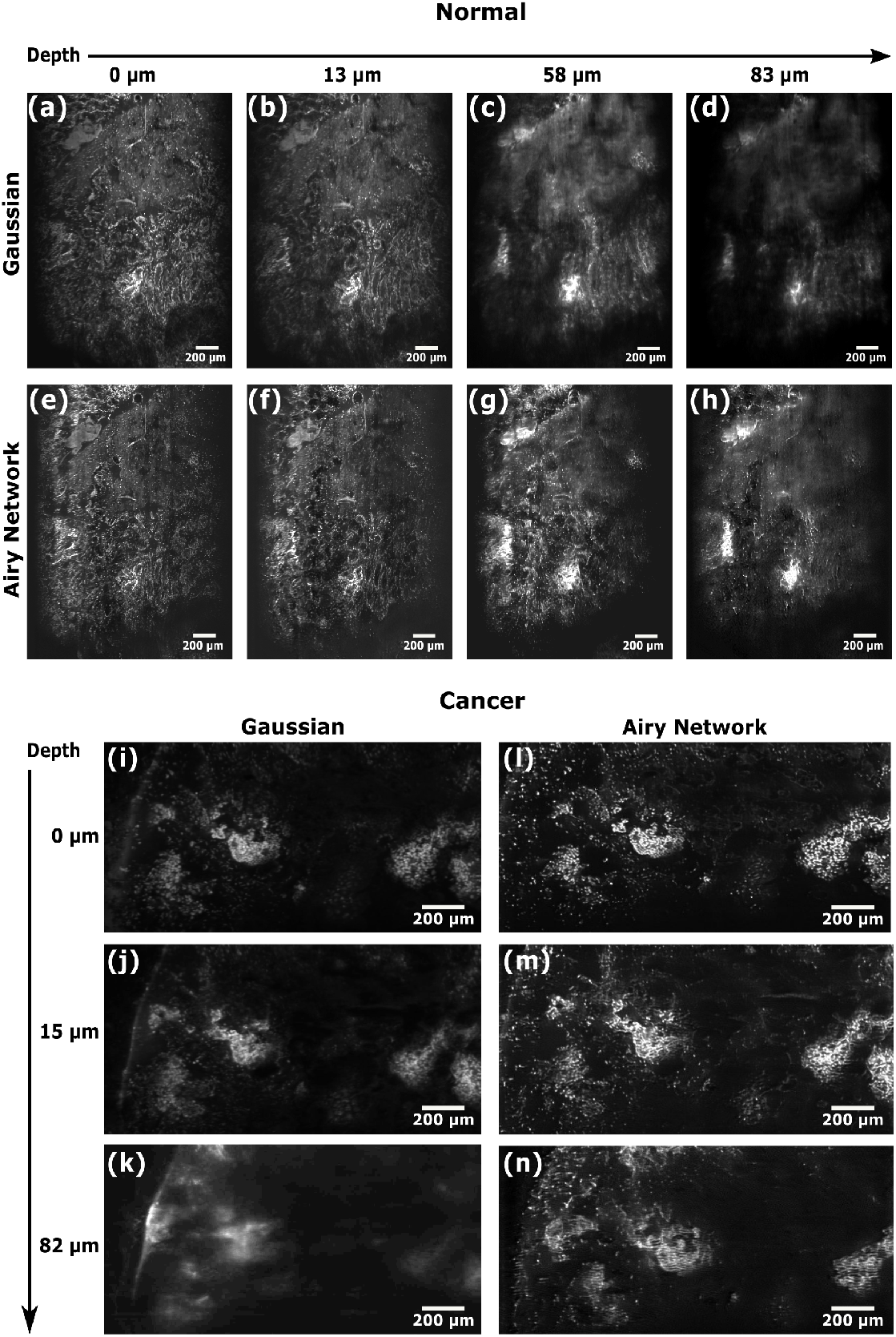
Images of healthy and cancerous colon tissue stained with acridine orange. The images of normal colon were acquired at different depths into the sample with both the (a–d) Gaussian and (e–h) Airy beam. The images of cancerous colon were also acquired at different depths into the sample with both the (i–k) Gaussian and (l–n) Airy beam. The Airy images are deconvolved and the deep learning algorithm is applied to them.

On the surface of colon cancer tissue (Fig. 5(i) and (l)), the crypts are still visible but there are also bright patches showing accumulations of nuclei, which are a common feature in cancer tissues. As for normal colon, the images acquired with the Airy beam to which the deep learning algorithm has been applied show better contrast with respect to those acquired with the Gaussian beam and allow single nuclei to be seen even at a depth of 82 *μ*m (Fig. 5(k) and (n)). This ability to follow structures in three dimensions, under the epithelial surface, may provide additional information on the direction and extent of cancer cell invasion.

### 3.5 Breast tissue

In the carcinoma of the breast, islands of epithelial cells within glandular and duct tissues undergo malignant transformation. Conventional treatments involve local surgical excision with or without hormone or chemotherapy. While imaging the tissue we have noticed that, compared to the healthy colon tissues which lay flat on the glass coverslip, the malignant breast tissue was substantially more heterogeneous, with an uneven surface topography that prevented uniform contact across the coverslip. The heterogeneity was particularly present in areas of tumour.

Figure 6 shows images of benign and malignant breast cancer tissues taken at the surface and at depth with the Gaussian light sheet and with the Airy light sheet to which the deep learning algorithm was applied. The images in Figs. 6(a-d) represent benign fibroadenoma with collagenous stroma and distorted, slit-like elongated ducts. The images in Figs. 6(e-h) represent invasive ductal carcinoma consisting of a broad sheet of densely packed tumour cells appearing as hyper-fluorescent dots. While for the colon tissue the difference between the Gaussian and deep-learning Airy images was more striking at depth, in the case of breast tissue the difference was visible even near the surface. In particular, the images taken with the Airy light sheet look substantially sharper. This could be addressed by the fact that is more challenging to focus the Gaussian beam with a shorter depth of focus on an uneven surface of the sample with poor contact to the cover slide. Also, the self healing properties of the Airy light field make it resilient to aberrations at the surface, in which refractive index mismatch due to air gaps is present. This aberration is evident in the insets in Fig. 6.

**Figure 6:**
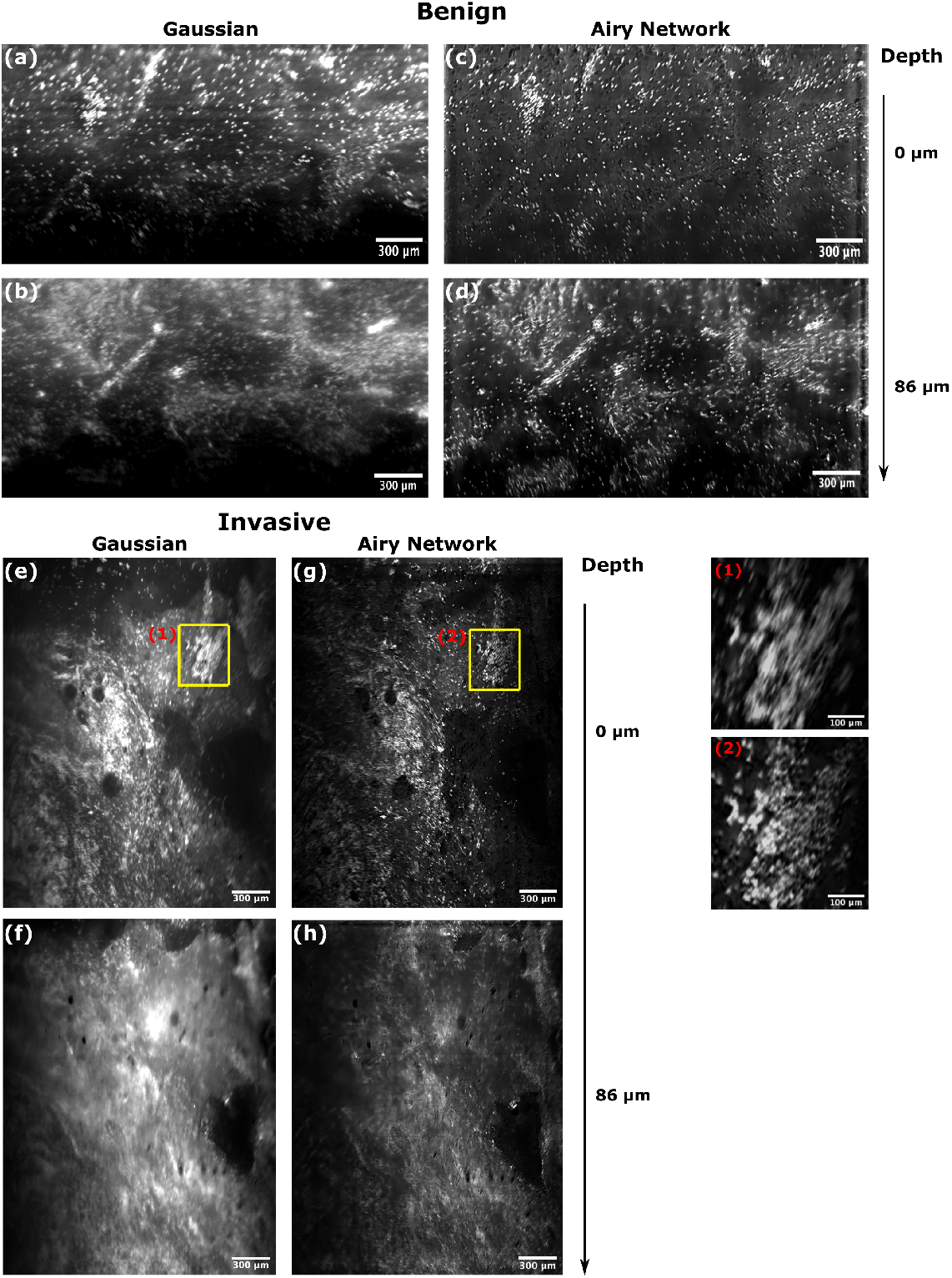
Images of benign breast tissue and invasive breast cancer tissue stained with acriflavine. The images were acquired at different depths into the sample with both the Gaussian and the Airy beam. The deep learning algorithm is applied to the Airy images after deconvolution. (a) and (c) show the surface of benign breast tissue acquired with the Gaussian and Airy beam, respectively. (b) and (d) show images of the benign tissue acquired at a depth of 86 *μ*m with the Gaussian and Airy beam, respectively. (e) and (g) show images of the surface of breast cancer tissue acquired with the Gaussian and Airy beam, respectively. Insets (1) and (2) show an agglomeration of nuclei from regions marked by a yellow rectangle in (e) and (g) respectively. (f) and (h) show images of the cancer tissue acquired at a depth of 86 *μ*m with the Gaussian and Airy beam, respectively.

## 4 Discussion

In this paper, we have developed an open-top light sheet microscope that combines the use of a non-diffracting Airy light field and a deep-learning method. This extends the field of view in depth to 1.6 mm and achieves a 3-mm wide field of view while recovering a sub-Nyquist-sampled resolution. The Airy light field demonstrated a higher axial resolution over a substantially larger depth range compared to the Gaussian beam. The advantages of the Airy light field have been demonstrated previously in [11]. The Airy beam was measured to improve the imaging depth through turbid tissues by approximately 30%. Further it was shown to improve contrast, with the largest benefits seen with increasing depth [19]. The improvements observed in this work agree with these observations. This system may further benefit from the addition of attenuation compensation to the Airy profile with a facile use of an additional exponential phase mask to concentrate a greater intensity of light at depth [26]. Additionally, we have incorporated Airy illumination in an open-top geometry, which greatly simplifies sample administration and positioning, especially for large tissue and heterogeneous sections with uneven surface topography. The SIL further removes the need for tissue embedding in agarose, or other refractive index-matching material. Such a geometry improves the prospects of technological adoption in medicine and biology.

The adoption of deep-learning in our system has demonstrated a near two-fold improvement in resolution whilst maintaining a wide field of view for light sheet imaging. This is consistent with previous observations in conventional widefield epifluorescence microscopy [18]. In LSM, deep learning has been applied previously in one instance, however, the training was performed using artificially simulated low-resolution images from high-resolution data [27]. In microscopy, deep-learning has been typically used to surpass the diffraction limit to achieve super-resolution microscopy [28]. However, in a broader context of image processing, deep-learning has been applied to a class of problems termed single-image super-resolution [29], the goal of which is to enhance resolution in poor sampling regimes. In this case, and as demonstrated in this paper, deep-learning may be used to effectively resolve sub-Nyquist-sampled features without necessarily breaching the fundamental diffraction limit. In this case, the network has to solve the challenge of effectively interpolating between sampled points based on known tissue features, rather than recovering information that cannot be fundamentally transferred by the optical system.

The use of deep-learning in microscopy has to be taken with care [28]. There are a variety of network models that can generate false features based on the features observed in the training data. The particular class of architecture used in this work is a method that relies on accurate pixel to pixel registration and non-linear pixel-wise mapping to the training data [18]. Thus, the model maybe conceptualised as an efficient non-linear deconvolution method based on a training set that captures typical microscopy aberrations. As described in the supplement to [18], the best results maybe obtained bytraining a new network with each individual imaging setup;however, we found that a good performance was achieved with a network pre-trained on widefield epifluorescence for both the Airy and Gaussian illuminations. Whilst the imaging modalities are different, many aberrations and losses in image quality remain consistent between methods, and are fundamental to all optical systems. Thus, due to the training with a broad range of samples and stains in [18], the pre-trained network demonstrated substantial and relevant improvements as seen in the LSM results presented in this paper. Using such a pre-trained network simplifies the technical requirements to adoption from multi-day training with a high-end workstation to ready sub-second reconstruction per image with a generic personal computer.

The research data underpinning this publication can be accessed at https://doi.Org/10.17630/7cee889f-aa36-4c27-a485-262c8a5d336b[30].

## 5 Conclusion

We have developed a light sheet microscope for widefield imaging of tissue volumes that incorporates an Airy light field to achieve imaging at an extended depth, and a deep-learning super-resolution architecture to achieve a two-fold improvement to the spatial dynamic range maintaining a wide field of view. We have demonstrated the promise of the system to enhance imaging of amyloid plaques in Alzheimer’s mouse brain and to distinguish healthy and cancerous colon and breast tissues from biopsies. The system has promise for rapid, facile, widefield and volumetric imaging of tissue sections in medicine and biology, and has prospects for simple adoption in other geometries of light-sheet microscopy.

## Acknowledgments

We thank Dr Adam K. Glaser for sharing the design of the SIL lens, and Dr Graham D. Bruce for a detailed review of the manuscript. We thank Imperial College Tissue Bank and Imperial College Experimental Cancer Medicine Centre (ECMC) for providing the breast tissue samples and Tayside Tissue Bank for providing the colon samples. Author Contributions: SC performed the main experimental studies with contributions from PBP. PW performed the numerical analyses. SS, KV and CSH provided samples. JN and FG assisted in the early stages of the project. SC, PW and KD wrote the paper which was reviewed and approved by all authors. KD initiated and supervised the study.

